# A NanoFE Simulation-based Surrogate Machine Learning Model to Predict Mechanical Functionality of Protein Networks from Live Confocal Imaging

**DOI:** 10.1101/2020.03.27.011239

**Authors:** Pouyan Asgharzadeh, Annette I. Birkhold, Zubin Triverdi, Bugra Özdemir, Ralf Reski, Oliver Röhrle

**Affiliations:** Institute for Modelling and Simulation of Biomechanical Systems, University of Stuttgart, Stuttgart, Germany; Stuttgart Center for Simulation Science (SC SimTech), Stuttgart, Germany; Plant Biotechnology, Faculty of Biology, University of Freiburg, Freiburg, Germany; Signalling Research Centres BIOSS and CIBSS, Freiburg, Germany; Cluster of Excellence livMatS @ FIT - Freiburg Centre for Interactive Materials and Bioinspired Technologies, Freiburg, Germany

**Author notes:** Corresponding author. Email address (Pouyan Asgharzadeh), URL: http://bit.ly/2Tqx_PA (Pouyan Asgharzadeh). These authors contributed equally to this work (Pouyan Asgharzadeh, Annette I. Birkhold). Currently an employee of Siemens Healthcare GmbH (Annette I. Birkhold).

**Keywords:** structure-function relationship, protein network, machine learning, finite element analysis, plastoskeleton

## Abstract

Sub-cellular mechanics plays a crucial role in a variety of biological functions and dysfunctions. Due to the strong structure-function relationship in cytoskeletal protein networks, light can be shed on their mechanical functionality by investigating their structures. Here, we present a data-driven approach employing a combination of confocal live imaging of fluorescent tagged protein networks, in-silico mechanical experiments and machine learning to investigate this relationship. Our designed image processing and nanoFE mechanical simulation framework resolves the structure and mechanical behaviour of cytoskeletal networks and the developed gradient boosting surrogate models link network structure to its functionality. In this study, for the first time, we perform mechanical simulations of Filamentous Temperature Sensitive Z (FtsZ) complex protein networks with close-to-reality network geometry depicting its skeletal functionality inside organelles (here, chloroplasts) of the moss *Physcomitrella patens*. Training on synthetically produced simulation data enables predicting the mechanical characteristics of FtsZ network mechanics purely based on its structural features (*R*^2^ ≥ 0.93), therefore allowing to extract structural principles enabling specific mechanical traits of FtsZ, such as load bearing and resistance to buckling failure in case of large network deformation. The presented method and the specific in silico findings from its application may allow in the future to reproduce mechanical cell responses in engineered environments.

## 1. Introduction

Bio-polymer networks are pervasive as key promoters of strength, support and integrity. This is true irrespectively of its scale, i.e., from the nano-scale of the cytoskeleton to the macro-scale of connective tissues. As cells sense external physical signals and translate them into a cellular responses, cellular mechanics has been proven to be crucial for a wide range of biological functions and dysfunctions. In particular cytoskeletal protein networks exhibit strong structure-function relationships, e.g., the role of microtubule network during mitosis [1], cell movement with the help of actin assembly/disassembly [2] or utilizing intermediate filament networks for stabilizing mechanical stresses [3]. In all these processes protein networks transmit besides biochemical also biophysical cues from the cell microenvironment that trigger and regulate cell behaviors. Investigating the structure of protein networks allows deeper insights into cellular functionality and dysfunctionality and may further elucidate many critical cell responses observed in vivo. Understanding these cellular responses in 3D may further lead to the development of functional and biomimetic materials for engineering the 3D cell microenvironment able to control cell behaviors in 3D and may advance the fields of tissue regeneration and in vitro tissue models.

In the recent decades, taking advantage of new methodological developments in experimental and computational physics and applying them to biological systems allowed substantial progress in elucidating particular mechanical phenomena to biological function. It have shown that mechanical processes convey biochemical signals and are therefore crucial for cell functions including proliferation, polarity, migration and differentiation. Further, connections between the mechanical properties of cells and initiation and pathological progression of cancer were established [4, 5]. Mendez et al. and Liu et al. showed that the epithelial-to-mensenchymal transition leading to cancer metastasis are linked to changes in mechanical characteristics of the cytoskeleton influencing the vimentin network [6, 7] as well as the polarity of the cell [8]. Moreover, cancer cells are typically found to be softer than normal cells. A decrease in the level of actin in the cytoskeleton of cancerous cells was linked to changes in the mechanical properties of the cell [9]. Such research underpins the importance of linking molecular changes within the cytoskeleton to structural and functional changes of the entire cell and therefore changes to the tissue.

In summary, in-depth knowledge of cellular and sub-cellular mechanics might allow the identification and classification of cells at different physiological and patho-physiological stages. However, to do so, new approaches need to be developed that are capable of simultaneously performing structural and mechanical analysis of sub-cellular structures in a (semi-)automated way. Mechanical stability and its contribution to shaping processes on the molecular scale are far from being completely understood. Further, it is not clear, if mechanical processes, besides conveying biochemical signals, also purely convey mechanical signals to invoke structural changes. The concept of the cytoskeleton as a shape-determining scaffold for the cell is well established [10], however, the tight coupling of actin, microtubule and intermediate filament networks impedes a separate analysis. To date, computer models of cytoskeletal biopolymer networks are based on models that represent the geometry in a (strongly) simplified way [11–15]. In depth analysis of structure-function relationships, however, require detailed structural and functional modelling.

Development of such models requires a protein network with similar structural functionality to cytoskeletal networks while being structurally less complicated, therefore allowing a detailed validation of the derived results. Proteins homologous to tubulin, which is part of the eukaryote cytoskeleton, such as the Filamentous Temperature Sensitive Z (FtsZ) protein family in the chloroplasts of the moss *Physcomitrella patens* are excellent examples for this purpose. FtsZ generates complex polymer networks, showing striking similarity to the cytoskeleton, and hence were named plastoskeleton [16]. In bacteria, FtsZ is a part of the bacterial cytoskeleton providing a scaffold for cell division [17–19]. Coassembly experiments provide evidence that FtsZ2 controls filament morphology and FtsZ1 promotes protofilament turnover. It is suggested, that *in vivo*, FtsZ2 forms the chloroplast Z-ring backbone while FtsZ1 facilitates Z-ring remodeling [20]. Moreover, as chloroplasts in loss-of-function mutants show distinct shape defects, FtsZ networks might provide scaffolds that ensure the stability and structural integrity of the chloroplast [21]. Additionally, gene knock-out experiments have shown that the FtsZ network is capable of undergoing large deformations upholding its structural integrity [22]. This adaptive stability is presumably linked to the developed structural characteristics of FtsZ network; making the cytoskeletal FtsZ network an ideal first application for introducing and testing a simulation-based method that aims at identifying a link between structural features of a cytoskeletal network and its mechanical functions.

State-of-the-art microscopy imaging techniques permit resolving micro-structural details of protein networks. Computational analysis of acquired images facilitates the quantification of components and its assembly to networks [23], and may allow tracking structural changes of the network assembly triggered by internal or external stimuli, i.e., connecting the structure to functionality or distinguishing between network types [24]. Machine learning (ML) algorithms have proven to be remarkably capable of automating such complex image analysis tasks [25] and of correlating image content to biological structural functionality [26–28]. Recently, the concept of ML-based surrogate models has proven to be highly advantageous in accelerating the performance of numerical simulations of complex mechanical environments [29] as well as predicting material properties [30]. A ML-based approach could link structural features to mechanical characteristics and would provide a way to answer questions like “How are FtsZ biopolymers capable of exhibiting adaptive stability?” or “Interplay of which structural changes in the cytoskeleton of a cancerous cell leads to adapting stiffness?”

To overcome the challenge of relating structure to function of cytoskeletal protein networks, we present an ML approach applied to 3D live laser scanning confocal microscopy images. The outcome is an end-to-end tool that links structural features associated with the cytoskeletal network type to its mechanical behaviour and therefore enables a fast evaluation of structure-function relations on the sub-cellular scale. This is carried out by combining an in silico mechanical characterization of protein networks through 3D nano finite element modeling and an automatic mapping of structural features to the mechanical network responses. The introduced method consists of two different models. The first model classifies the respective protein networks based on their structural features by exploiting an image processing and a gradient boosting classification model. The second one creates an in silico surrogate model to predict the sub-cellular mechanical responses of the network. Analyzing the prediction process of the surrogate model based on the structural feature allows us to deduct the presumed structure-function relationship. The method is tested and applied to elucidate isoform-specific structure-function relationships of FtsZ networks.

## 2. Materials and Methods

### 2.1. Materials

The “Gransden 2004” ecotype of the moss *Physcomitrella patens* ((Hedw.) Bruch & Schimp., IMSC accession number 40001) was cultivated in bioreactors [31].

### 2.2. Molecular Biology and Moss Transfection

RNA isolation, molecular cloning and moss transfection were described previously in detail [23, 24] and are therefore given here only in a shortened version. Total RNA was isolated from wild type *Physcomitrella patens* protonema using TRIzol Reagent (Thermo Fisher Scientific) and used for cDNA synthesis using Superscript III reverse transcriptase (Life Technologies, Carlsbad, CA, USA). The coding sequences of PpFtsZ1-2 and PpFtsZ2-1 were PCR-amplified from this cDNA and cloned into the reporter plasmid pAct5::Linker:EGFP-MAV4 (modified from [32]) to generate the fusion constructs PpAct5::PpFtsZ1-2::linker::EGFP-MAV4 and PpAct5::PpFtsZ2-1::linker::EGFP-MAV4. Moss protoplasts were isolated and transfected with 50 *µg* of each of these plasmids, according to the protocol described by Hohe et al. [33]. The transfected protoplasts were incubated for 24 h in the dark, subsequently being returned to normal conditions (25.1°*C*; light-dark regime of 16 : 8*h* light flux of 55 *µmol s*^−1^*m*^−2^ from fluorescent tubes, Philips TL - 19 - 65W/25).

### 2.3. Laser Scanning Confocal Microscopy Imaging

In 4 − 7 days after transfection, the protoplasts were concentrated to a volume of 100 *µl*, and 20 *µl* of this protoplast suspension was used for imaging. Confocal laser scanning microscopy (CLSM) images were taken with a Leica TCS SP8 microscope (Leica Microsystems, Wetzlar, Germany), using the HCX PL APO 100x/1.40 oil objective and applying the microscopy conditions described previously [23, 24]. A selection of images visualising FtsZ networks is depicted in Fig. 1a. To summarise, the zoom factor was 10.6, the voxel sizes were 21 *nm* in the *X* − *Y* dimensions and 240 *nm* in the *Z* dimension and the pinhole was adjusted to 0.70*AU* (66.8 *µm*). For the excitation, a WLL laser was applied at 488 *nm* with an intensity of 4%. The detection ranges were set to 503−552 *nm* for the EGFP fluorescence and 664 − 725 *nm* for the chlorophyll autofluorescence. All images were deconvolved using Huygens Professional version 17.04 (Scientific Volume Imaging, The Netherlands). This resulted in a dataset of *n* = 37 3D CLSM images (i.e., 21 FtsZ2-1 and 16 FtsZ1-2 isoforms).

**Figure 1:**
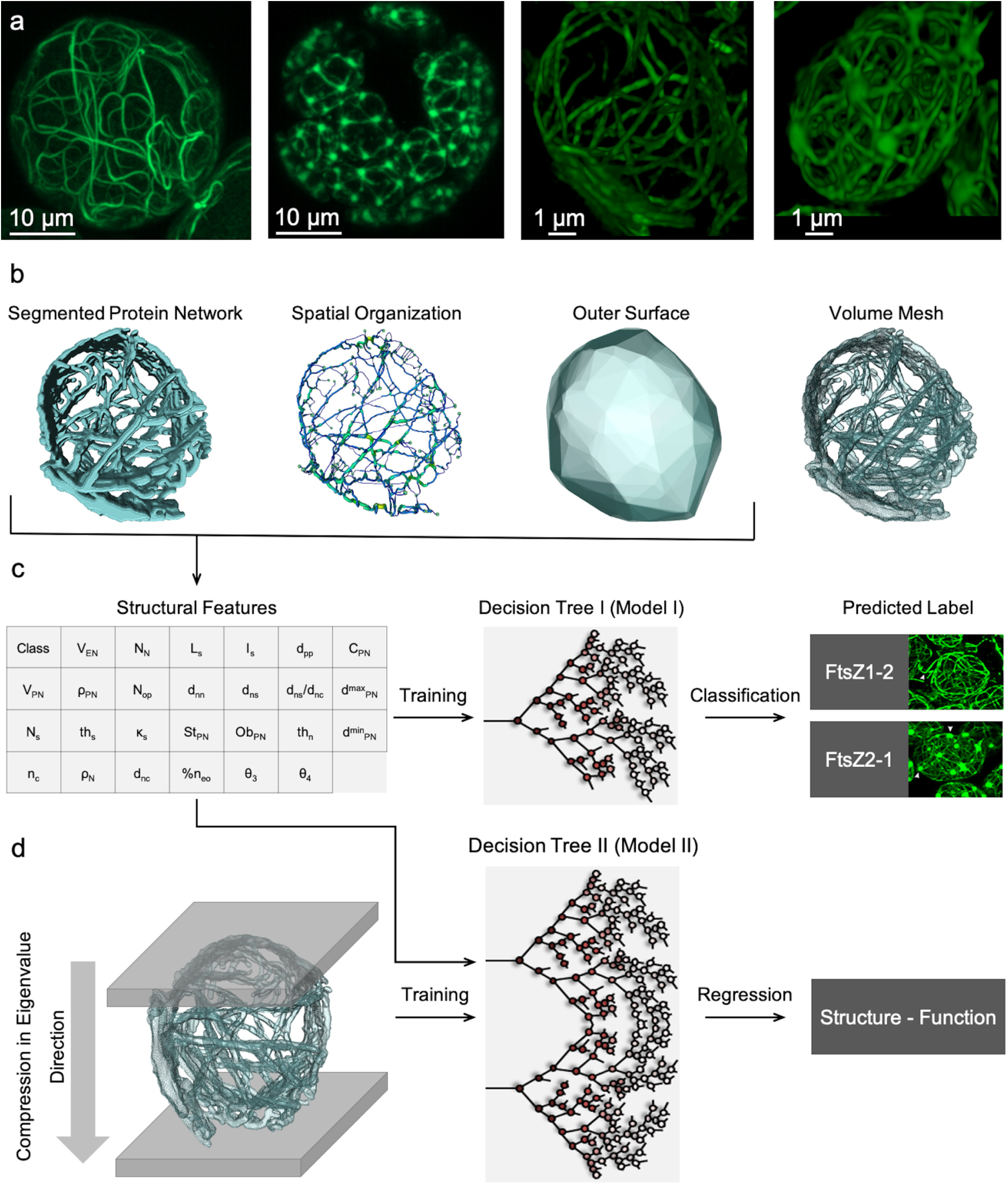
Correlating structure of FtsZ network to its mechanical functionality utilizing CLSM images. a) Sample 3D CLSM images of FtsZ1-2 and FtsZ2-1 networks in a cell (two left images) and single FtsZ1-2 and 2-1 networks (two right images), respectively. b) Sample of a 3D segmented image and its spatial graph, convex hull and mesh. c) The 26 shape and element descriptors that are extracted and used as input features to train a gradient boosting model for classifying FtsZ1-2 and FtsZ2-1 isoforms. d) A second gradient boosting model (regression) is trained on the structural features to predict the results of the mechanical simulation of compressing the network in its principal directions (3 Eigenvectors determined from segmented images).

### 2.4. Image Processing to Extract Structural Features

A set of 26 structural features describing the assembly of protein networks from global and local perspectives is extracted from each network. Here, only a short description of the workflow steps and features are given, details as well as a validation have been reported previously [23].

#### 2.4.1. Image Pre-processing

Images are segmented using an adaptive local threshold, 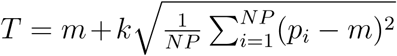, with *NP* = 10 ∗ 10 ∗ 10 being the local window size, *m* is the average pixel intensity in the window, *p*_*i*_ denotes the intensity of pixel *i* and *k* = 10 as a constant value. Next, by calculating the convex hull of the segmented network a solid outer surface representing the volume enclosing the network is determined. To extract the structural features of the network, a transformation to a spatial graph consisting of points, nodes and segments is performed. This transformation consists of following steps: 1. determining the centerline of each filament based on calculating a distance map for each foreground voxel from the edge voxels, and 2. placing points at the centerline of the filaments where either thickness or the direction of the filament changes. The resulting hierarchy of structural elements of the spatial graph reads as: 1. points, 2. elements as the connection between points, 3. nodes, as points that are connected to more than two other points, 4. segments as summation of elements from one node to another (filaments), 5. connections as the meeting points of filaments in a node. This numerical representative in form of a spatial graph allows determining structural features (Fig. 1b).

#### 2.4.2. Features Describing 3D Network Morphology

Seven shape descriptors, which are extracted by analyzing the segmented network and its convex hull, quantitatively describe the morphology of the network [23]. These features consist of: 1. enclosed volume of the network, *V*_*EN*_, as the volume of the convex hull, 2. network volume, *V*_*PN*_, as the volume of the segmented protein network, and 3. the network volume density, *ρ*_*PN*_, as the ratio of enclosed volume to network volume. Furthermore, a shape matrix representing the covariance of the convex hull is calculated. Building upon the eigenvalues of the shape matrix, the 4. greatest and 5. smallest diameters of the network, 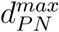 and 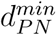. 6. stretch of the network, *St*_*PN*_, and 7. oblateness of the network, *Ob*_*PN*_ are determined.

#### 2.4.3. Features Describing 3D Network Structure

19 local structural features calculated from the spatial graph of the network consist of nodal features (1. number of nodes, *N*_*n*_, 2. thickness of nodes, *th*_*n*_, 3. node density, *ρ*_*n*_, 4. node-to-node distance, *d*_*nn*_, 6. node-to-surface distance, *d*_*ns*_, 6. node-to-centre distance, *d*_*nc*_, 7. number of open nodes, *N*_*op*_. 8. compactness, *C*_*PN*_, defined as *C*_*PN*_ = (*d*_*nc*_ − *d*_*ns*_)*/d*_*nc*_. 9. node-to-surface to node-to-center ratio), segment features (10. number of segments, *N*_*s*_, 11. segment length, *L*_*s*_, 12. segment curvature, *κ*_*s*_, 13. mean segment thickness, *th*_*s*_, 14. segment inhomogeneity, *I*_*s*_, 15. segment point-to-point distance, *d*_*pp*_), and connection feature (16. mean number of connections per node, *n*_*c*_ 17. percentage of open nodes, *n*_*oe*_, 18./19. mean angles between segments in a connection with 3/4 filaments meeting, *θ*_3_ and *θ*_4_). The morphological and structural features are calculated by a set of in-house Matlab codes (Matlab 2019a, MathWorks, USA).

### 2.5. Mechanical nano-FE modeling

To investigate the mechanical response of the protein networks to external load, we designed a generic in-silico experiment reflecting a compression against a plate, hence, a scenario that is typically used to experimentally investigate the mechanical behaviour of whole cells [34]. To capture the overall mechanical behavior of each network in a comparative manner, compression tests along the three principal axis of each system were modeled employing a nano-FE approach. All simulations were done with the finite element analysis software Abaqus 6.14 (Dassault Systèmes, France). It is important to note that, the goal of this setup is to depict the mechanical behavior of the protein network morphology rather than replicating the real physical condition and dynamics of the biopolymers in their biological roles. Such a task would require consideration of highly complicated interactions of the network with its surrounding, which are not completely understood to date.

#### 2.5.1. 3D Protein Network Model Generation

For all samples, protein network surface meshes were defined from the segmented images using a triangular approximation algorithm coupled with a best isotropic vertex placement algorithm to achieve high triangulation quality [35]. The surface area of the resulting surface mesh was calculated and further remeshed using *n*_*t*_ = *ρ*_*t*_*A*_*t*_ triangles for the remeshed surface, where *ρ*_*t*_ = 900 [*triangles/µm*^2^] is the constant surface mesh density and *A*_*t*_ denotes the surface area. Furthermore, the remeshed surface was smoothed by shifting the vertices towards the average position of its neighbours. The enclosed surfaces were filled with volumetric tetrahedral elements, resulting in an adaptive multi-resolution grid (Fig. 1b) using FEI Amira 6.3.0 (Thermo Fisher Scientific, USA).

The principal directions of a network were determined based on its convex hull and shape matrix. The eigenvectors of the shape matrix (*EV* 1, *EV* 2 and *EV* 3), which are orthogonal to each other, represent the network’s principal directions, *V*_*i*_ (*i* = 1, 2, 3). The mesh is then transformed to the coordinate system spanned by *V*_*i*_. Afterwards, along each *V*_*i*_, the pair of nodes exhibiting the largest distance in-between the two points and in the direction of *V*_*i*_ were determined and named *N*_*i*1_ and *N*_*i*2_ and *i* = 1, 2, 3, respectively.

For each considered protein network, the geometry of the protein network was imported to Abaqus and three compression simulations (one per direction *d*) were carried out. For each simulation, the initial set-up was determined by first identifying the initial position of two parallel rigid plates, which are defined for each simulation in direction d by its normal vector 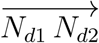 (with *d* = 1, 2, or 3), and the respective nodal points *N*_*d*1_ and *N*_*d*2_ (cf. Fig. 2a-c).

**Figure 2:**
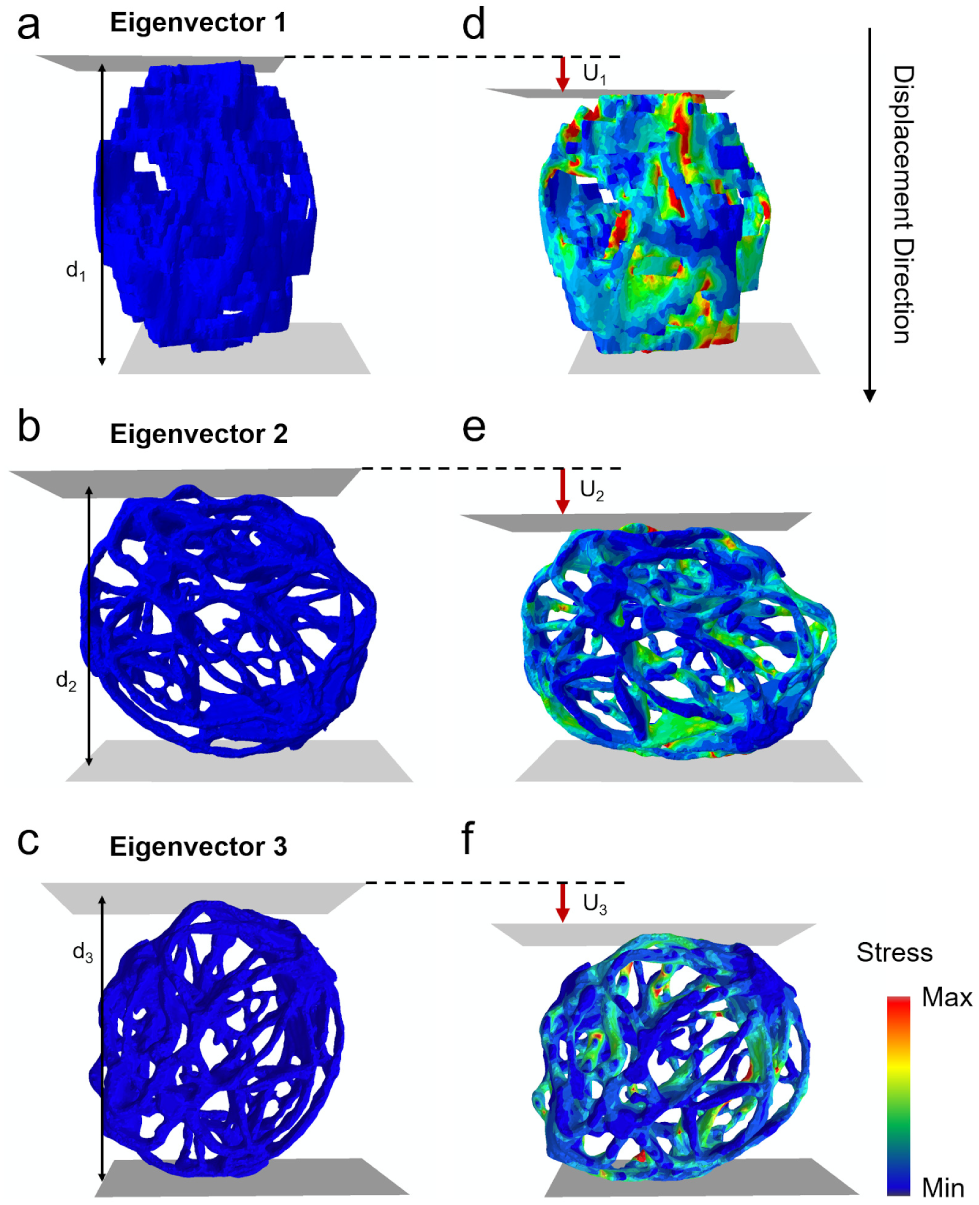
Simulation setups. a-c) Initial set-up for virtual compression experiments of a sample protein network in primary directions EV1, EV2 and EV3, respectively. d-f) Stress distribution after applying a displacement (*α* = 0.02) to the upper plate in EV1, EV2 and EV3 directions, respectively. The displacements are scaled by a factor of 5.

#### 2.5.2. Governing equations

The simulations were carried out by solving the balance of linear momentum,

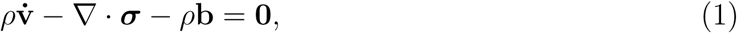

in an explicit manner, where *ρ* is the mass density, **v** denotes the velocity, ***σ*** describes the Cauchy stress, and **b** are the body forces. The contact between the protein network and the rigid plates was chosen as a rough contact meaning that any two points, which come in contact, will stick together (relative penetration tolerance: 1 e−3). The governing equation is discretized using the FE method, tetrahedral elements and linear spatial Ansatz functions.

#### 2.5.3. Boundary Conditions

The generic boundary conditions for each simulation setup (one simulation for each primary directions: *EV* 1, *EV* 2 and *EV* 3) consist of applying displacement boundary conditions at node *N*_*d*1_ to mimic compression experiments. The displacement itself is applied in the 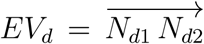 direction and in fractions (*α*) of the initial distance, 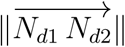, between the two plates (cf. Fig. 2). Therefore, the amount of displacement along the respective Eigenvector *EV*_1_ is defined by 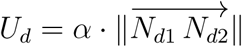.

To investigate anisotropy in the mechanical response of the network, compression tests along all three primary directions were performed and compared. Furthermore, for analyzing changes in the structural behavior with increasing deformation grade, we increased the displacement of the plate gradually in steps of *δα* = 0.02. We applied a total of 10 steps, which is equivalent to *α* = 0.20. Due to no apparent significant differences in the mechanical behaviour of the network between the three directions at *α* = 0.02 and the required computational resources, we chose to focus only on one direction to continue the simulations for *α* = 0.02 → 0.20.

#### 2.5.4. Constitutive Law and Material Parameters

Employing the concept of linear elasticity, the stress tensor ***σ*** is given by

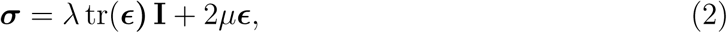

where ***ϵ*** denotes the strain tensor and ***I*** is the second-order identity tensor. Further, λ and *µ* are the first and the second Lamé coefficients, respectively. The Lamé coefficients are related to Young’s modulus *E* and Poisson’s ratio *v* by

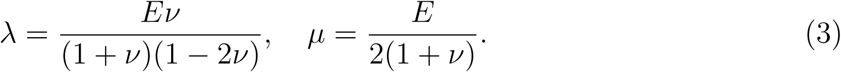

In classical continuum mechanics, the material parameters (here λ and *µ* or *E* and *ν*) are obtained by making a constitutive assumption, i.e., selecting a particular phenomenological constitutive law (here the form of ***σ***) and ensuring that the computed stresses match the experimental ones. The mechanical behavior of filamentous biopolymers is, however, commonly quantified by means of the flexural rigidity, *κ* [36–38], *which is the force couple required for one unit of curvature [39]. The flexural rigidity, κ*, is defined as *κ* = *EI*, where *I* is the second moment of inertia. In the context of protein filament mechanics, the flexural rigidity is calculated as *κ* = *k*_*B*_*T l*_*p*_, where *k*_*B*_ = 1.38 × 10^−23^ J*/*K is the Boltzmann constant, *T* defines the temperature, and *l*_*p*_ denotes the corresponding thermal persistence length. Recently, persistence length and flexural rigidity of FtsZ filaments reported by Turner et al. as *κ* = 4.7 ± 1.0 × 10^−27^ Nm^2^ and *l*_*p*_ = 1.15 ± 0.25 *µ*m are commonly employed [40, 41]. The average thickness of filamentous elements of the FtsZ network has been reported to be 117 ± 28 nm [24]. Assuming circular cross sections, *I* equals 1.81 × 10^−29^ m^4^ [42]. Based on these values, we set within our simulations the elasticity modulus to *E* = 2.6 × 10^2^ Pa and the Poisson’s ratio to *ν* = 0.5, i.e., assuming incompressibility [43–45].

#### 2.5.5. Calculated Mechanical Parameters

We performed a total of 111 simulations (3 simulations per network) on a CPU cluster with 32 cores (4 AMD Opteron Socket G34 Eight-Core 6328, 3.2 GHz, 8C, Abu Dhabi). One simulation took on average 19 ± 7 hours (for the entire 20% compression). To quantitatively assess the mechanical behavior of the protein networks, we determined the mean stress and strain of the networks, 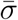 and 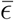, respectively, as the average of the L1 norms of the von Mises stresses and principal strains.

Cytoskeletal structures are reported to fail by buckling or rupture [46]. We therefore, further analyzed the structural stability of the network by calculating a buckling failure factor based on critical stresses *σ*_*crit*_ and a rupture failure factor based on critical strains *E*_*crit*_. Buckling of a single filament is assumed to occur if local von Mises stresses exceed a critical value. A filament is assumed to rupture, if strains locally exceed a critical strain value. To our knowledge, *σ*_*crit*_ and *ϵ*_*crit*_ of FtsZ have not been measured to date. However, despite fundamental structural differences, F-actin and FtsZ show similar mechanical behavior. The rigidity of F-actin is assumed to be *κ* = 7.5 · 10^−26^ Nm^2^ [36, 47] whereas the rigidity, which we assume for FtsZ filaments, is *κ* = 4.7 ± 1.0 × 10^−27^ Nm^2^ (*l*_*p*_ of F-actin: 1.77 *µm* and *l*_*p*_ of FtsZ: 1.15 *µm* [36, 40]). *Therefore, we use the values reported for F-actin (σ*_*crit*_ = 3.2 *Pa* and *ϵ*_*crit*_ = 0.2 [44, 47]). Since a local failure might not lead to a collapse of the whole network structure, we define failure factors based on the assumption that if a certain portion *m* of all elements 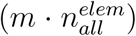 exhibit stresses or strains above the critical stress or strain value, the whole structure will fail by buckling or rupture of an individual or several segments, as shown for other biological materials [34, 48]. Since for protein networks, these threshold values have not been experimentally investigated yet, we report only the portion of elements that exceed a particular critical stress or strain value, i.e., the higher the values the higher the failure probability. We define the buckling failure factor as 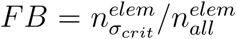, and the rupture failure factor as 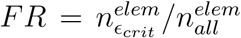, where 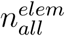 is the total number of elements, 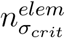 and 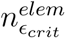 are the number of elements with stress and strain exceeding the critical buckling or rupture values, respectively.

### 2.6. Data-driven Analysis using Machine Learning

To relate the mechanical functionality of protein networks to their structure, we utilize a ML approach. To do so, we trained two sets of ML models on the dataset containing the 26 calculated structural features of the *n* = 37 protein networks. The aim of these ML models is to 1) perform a classification of the networks as well as an analysis of the structural features dominating the decision process and 2) map the structural features of the network to its mechanical behavior by employing a regression model. This further allows us to identify the most dominant structural features contributing to specific mechanical traits of the network.

#### 2.6.1. Classification of FtsZ Isoforms

We designed and trained a gradient boosting model [49], to perform the prediction task based on the extracted features (Fig. 1c):

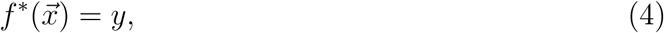

where we produce a prediction model (*f* ^∗^) in the form of an ensemble of weak prediction models to map the set of protein structural features:

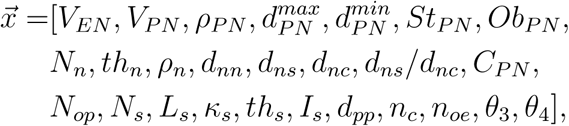

to an isoform (*y* : FtsZ1-2 or FtsZ2-1). During learning, we consecutively fit new models to provide a more accurate prediction [50], to construct the new models to be maximally correlated with the negative gradient of the loss function Ψ, associated with how wrong the prediction is. Given *N*_*n*_ training examples: 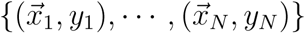, where 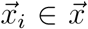 and *y*_*i*_ ∈ *y*, the gradient boosting decision tree model estimates the function *f* of future 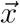 by the linear combination of individual decision trees

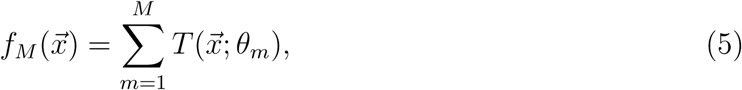

where 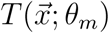 denotes the i-th decision tree, *θ*_*m*_ is its parameter set, *M* is is the number of decision trees. The final estimation is determined in a forward stage-wise fashion, i.e. based on an initial model 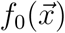 of 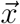, the model of step m is determined as:

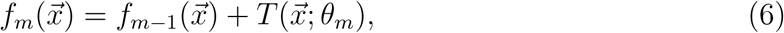

where 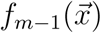 is the model in step *m* − 1. *θ*_*m*_ is learned by empirical loss minimization as

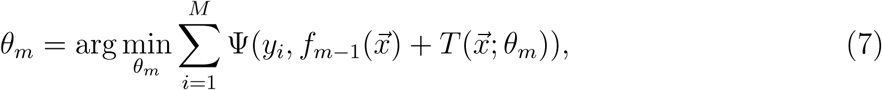

with the loss function Ψ. The assumption of linear additivity of the base function, leads to the estimation of *θ*_*m*_ for best fitting the residual 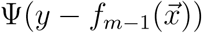. To this end, the negative gradient of the loss function at *f*_*m*−1_ is used to approximate the residual *R*:

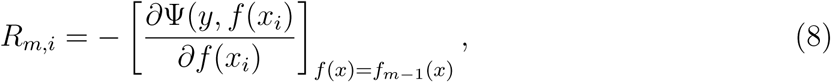

with i as the index of the i-th example.

We randomly divided the dataset into 80% training set (*n* = 30) and 20% test set (*n* = 7) with test set being stratified for number of isoforms. The model is trained on the training set employing a 5-fold cross validation [51] and applied on the test set.

#### 2.6.2. Surrogate Mechanical Model for Predicting Function from Structure

To investigate the structural approach(es) employed by nature to provide networks with specific mechanical functionality, e.g. adaptive stability, a set of surrogate models (SM) in the form of regression boosted gradient models are designed as:

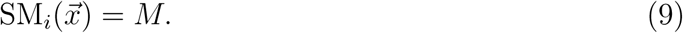

These 8 surrogate models serve as a tool to map the structural features (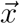, including the isoform class) of the protein networks to their mechanical behavior (Fig. 1d), with 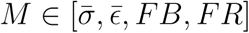 as the calculated mechanical parameter in *EV* 3 direction (cf. 2.5.5) for small and large deformations (*α* = 0.02 and 0.20) separately. The design and training of the regression surrogate models are similar to the described classifier model with the only difference being the definition of the loss function.

Here, the dataset is randomly divided into 90% training set (*n* = 33) and 10% test set (*n* = 4) with test set being stratified for number of isoforms. The model is trained on the training set employing a 5-fold cross validation and applied on the test set.

#### 2.6.3. Analyzing Feature Importance

We determined the importance of each structural feature in both, the classifier and the surrogate mechanical models. To do so, each feature is noised up and the plurality of out-of-bag vote and the reality are determined allowing to measure a wrong prediction rate [52] for each feature.

### 2.7. Statistical Analysis and Model Performance Evaluation

To distinguish the mechanical behavior of FtsZ1-2 isoforms from FtsZ2-1 isoforms, statistical analysis of 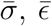, *FB* and *FR* was performed using repeated measures ANOVA and paired or unpaired student’s t-tests, as appropriate, followed by Bonferroni corrections for multiple comparisons. All values are presented as mean ± standard deviation and statistical significance was set to *p* < 0.05.

The performance of the both classifer model and the surrogate models is assessed by applying the models on the corresponding test set. The classifier model is evaluated by calculating the *F*1-score and the accuracy of prediction [53]. The performance of each surrogate models is assessed by calculating *R*^2^-values between the model predictions and simulation results. To further measure the differences between the surrogate model predictions and the true values of simulation results, we calculated the mean percentage error (*MPE*) of prediction as well as the slope of the linear fit for scattered data of model predictions vs. simulation results (*ŷ*).

## 3. Results

### 3.1. Effect of Load Direction on Mechanical Response

*N* = 16 FtsZ1-2 and *n* = 21 FtsZ2-1 isoforms images (see examples in Figure 3a, e) were processed. For each protein network, the image processing resulted in distinct spatial graphs (Fig. 3b, f), convex hulls (Fig. 3c, g; supplementary video 1), and FE meshes (Fig. 3d, h).

**Figure 3:**
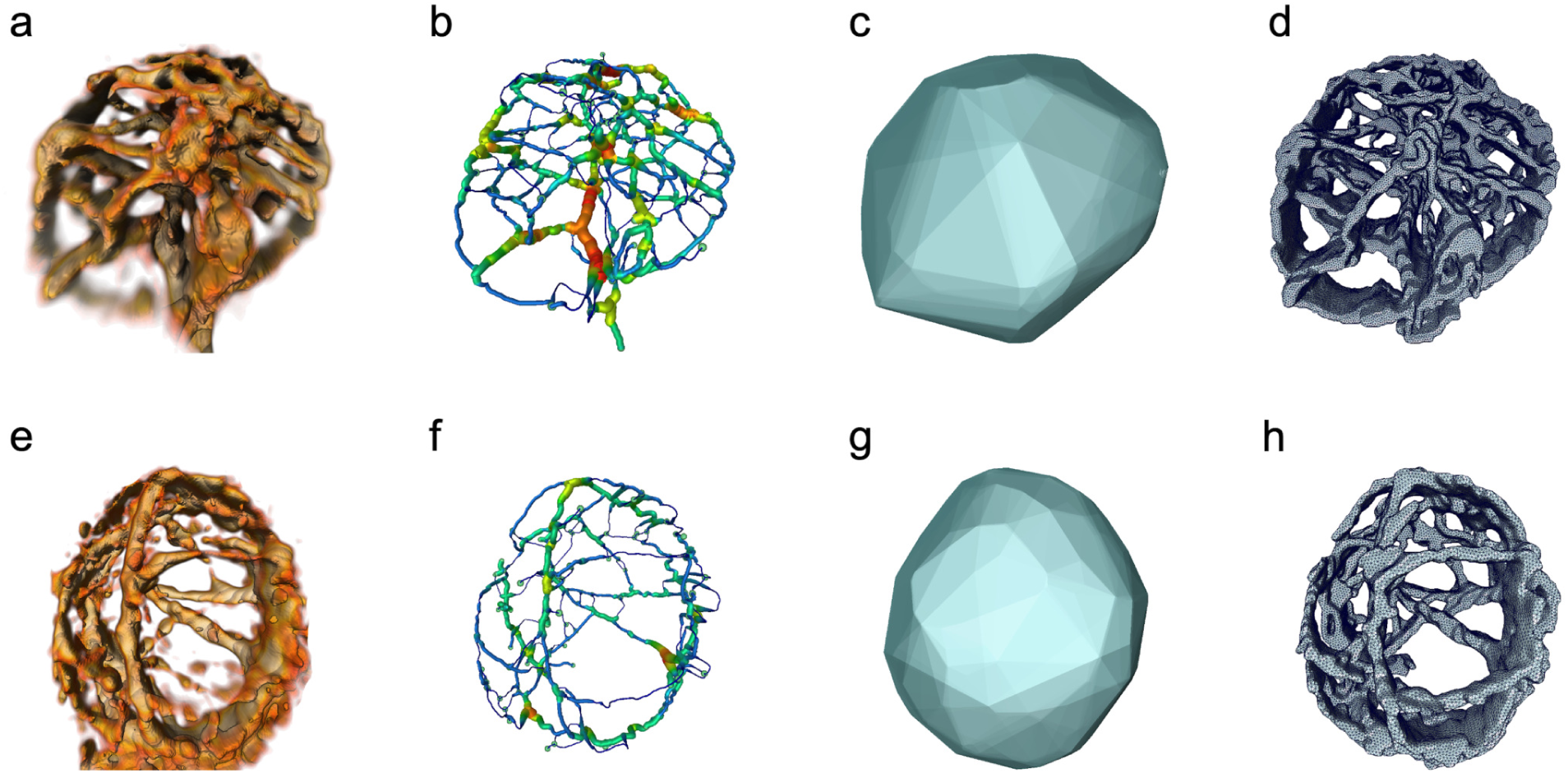
Image pre-processing of FtsZ1-2 (a-d) and FtsZ2-1 (e-h) isoforms. a) Sample 3D CLSM image of FtsZ1-2 isoform, b) resulting spatial graph, c) resulting convex hull and d) resulting volume mesh. e) Sample 3D CLSM image of FtsZ2-1 isoform, f) resulting spatial graph, g) resulting convex hull and h) resulting volume mesh.

All mechanical parameters (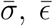, *FR* and *FB*) were affected by load direction as well as isoform type (ANOVA, *p* < 0.01). 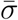, in the FtsZ1-2 isoform, was significantly lower for the EV2 load case than for the other two load cases (*p* = 0.01, Fig. 4a). In FtsZ2-1 bucking failure (*FR*) was significantly lower for the EV1 load case than the EV3 load case (*p* < 0.01, Fig 4d). Comparing the mechanical parameters (*x*, 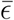, *FR, FB*) between the isoforms revealed that all mechanical parameters of the FtsZ2-1 isoform were for the EV2 loading case significantly higher than for the FtsZ1-2 isoform (*p* ≤ 0.04; Fig. 4a-d). Additionally, FtsZ2-1 responded to compression in EV3 direction with a significant higher 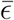 than FtsZ1-2 (*p* = 0.049; Fig. 4b).

**Figure 4:**
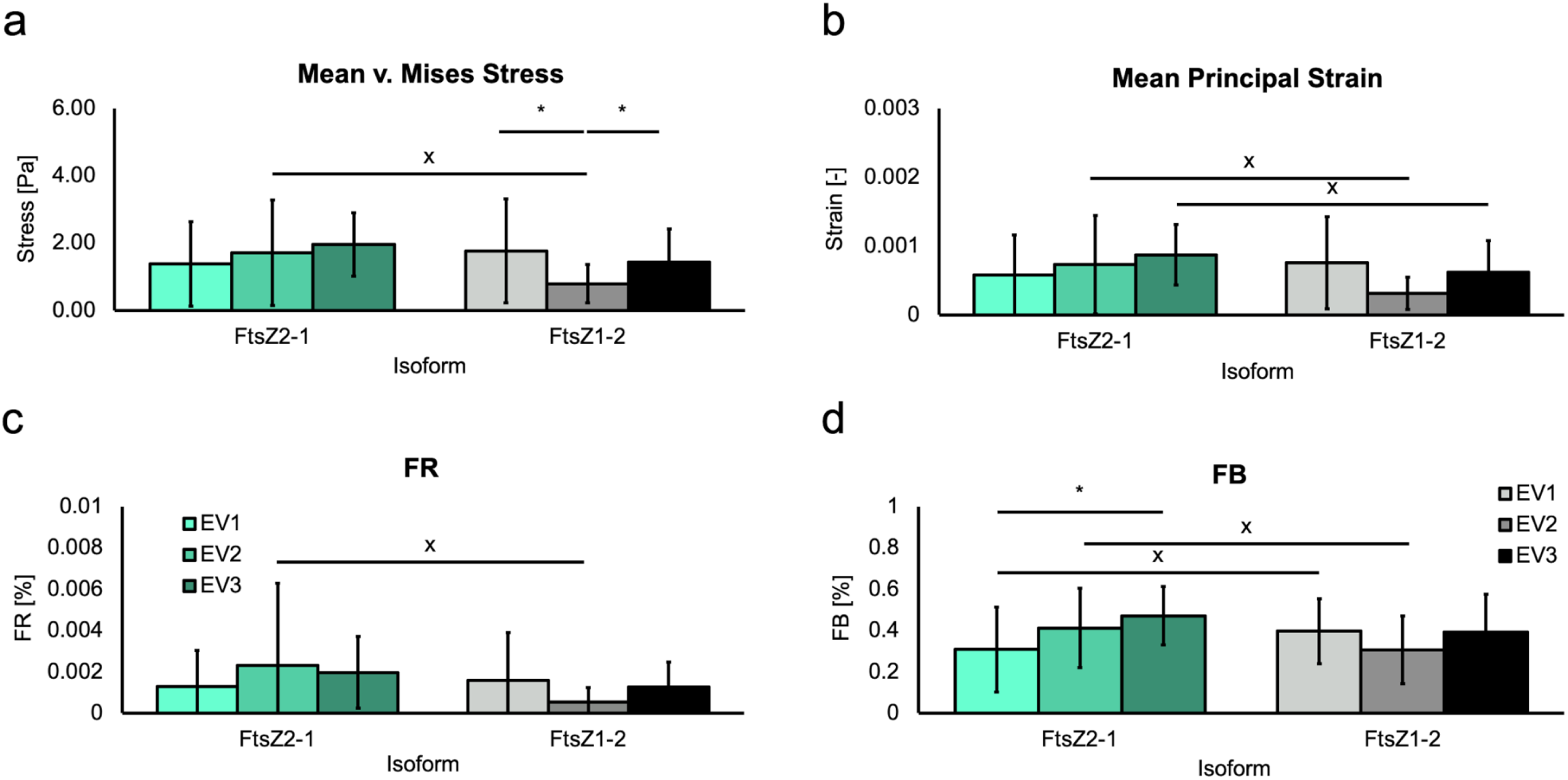
Mechanical responses to small deformations (2% compression). a) 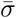. b) 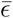. c) *FR* d). *FB*. Data is shown as mean±standard deviation. * denotes a significant difference between load directions (student’s t-test, Bonferroni correction), X denotes a significant difference between isoforms. Data is shown as mean±standard deviation.

### 3.2. Effect of displacement on mechanical response

The mechanical response with increasing compression was investigated only in EV3 direction. Increasing the compression of the isoforms from 2% to 20% (Fig. 5; supplementary video 2) revealed that, at all displacement steps, no significant difference between the two isoforms in the four calculated mechanical parameters occurred (Fig. 5c-f). For mean stress, mean strain and rupture failure factor (Fig. 5c-e), a gradual increase in both network types was detected with increasing displacements. In contrast, with increasing compression, *FB* converges toward a buckling failure factor of 1% (Fig. 5f).

**Figure 5:**
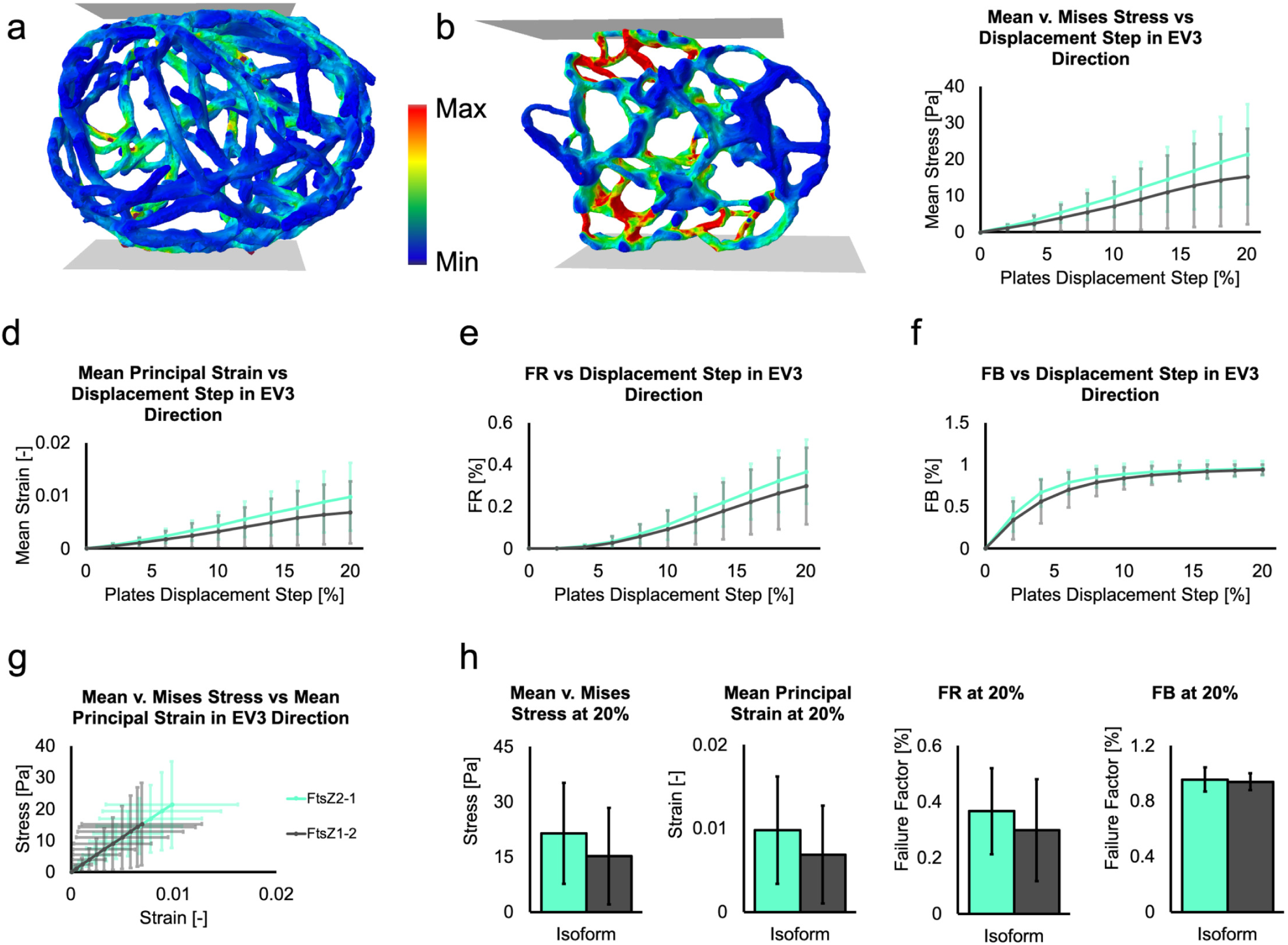
Changes in mechanical response with increasing compression. a, b) Stress distributions at 20% displacement in sample networks of FtsZ1-2 an FtsZ2-1, respectively. c) 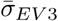. d) 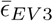. e) *FR*_*EV* 3_. f) *FB*_*EV* 3_. g) Mean stress vs mean strain in EV3 direction. h) Calculated mechanical parameters (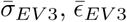 and *FB*_*EV* 3_ respectively) at the 20% displacement step. Results are presented as mean±standard deviations. Displacement step size is 2% with minimum displacement of 2% and maximum displacement of 20%.

Comparing the mechanical responses at 20% displacement shows no significant difference between the two FtsZ isoforms (Fig. 5h). At 20% displacement, the buckling failure factor (FtsZ1-2: 1.0±0.1% and FtsZ2-1: 1.0±0.1%) is significantly higher than the rupture failure factor (FtsZ1-2: 0.3±0.2% and FtsZ2-1: 0.4±0.2%, *p* ≤ 0.01). However, the first derivative of the failure factors with respect to the displacement (*FR*: FtsZ1-2: 0.04, FtsZ2-1: 0.05 and *FB*: FtsZ1-2: 0.00, FtsZ2-1: 0.00) shows that with increasing displacement, *FR* would presumably become the dominating failure factor.

### 3.3. Isoform Classification based on Structural Features

The classifier model trained on the 26 structural features reached 6 out of 7 correct predictions accuracy with an *F* 1-score of 0.88 on the test set. A correctly classified FtsZ2-1 isoform and a correctly classified FtsZ1-2 isoform, as well as the wrongly classified isoform (FtsZ1-2) are depicted in Fig. 6a-c, respectively.

**Figure 6:**
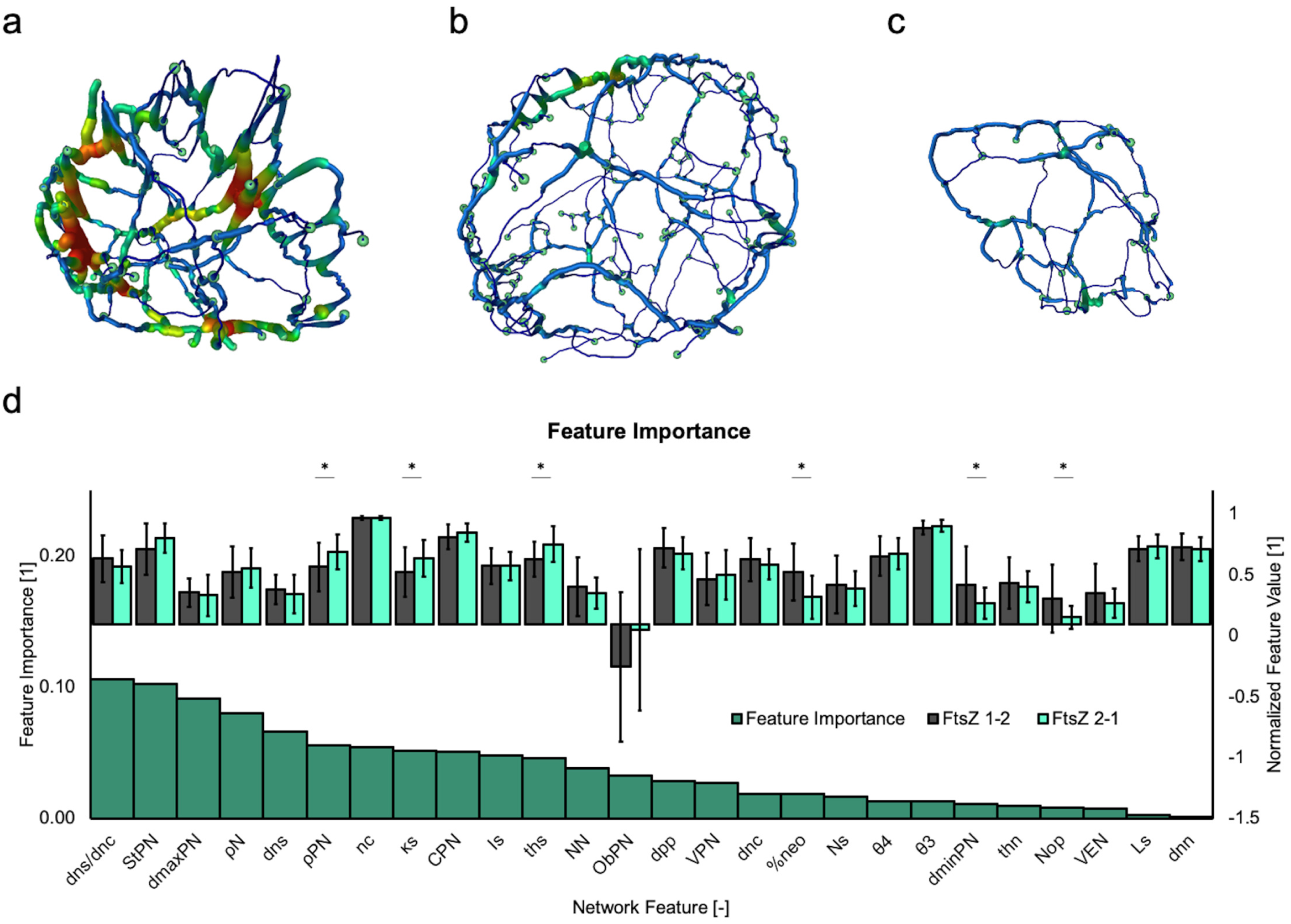
Classification of isoforms. a) A sample spatial graph of correctly classified FtsZ2-1. b) A sample spatial graph of correctly classified FtsZ1-2. c) A sample spatial graph of wrongly classified FtsZ1-2. d) Feature importance for the classification model as well as normalized feature values (normalized to maximum of each feature). Data shown as mean ± standard deviation. * indicates a significant difference between isoform (un-paired students’ t-test).

Analyzing the importance of each of the structural features in the classification model reveals which of the features contribute most and which least in terms of classifying isoform-inherent structural properties. The five most important structural features are the node-surface to node-center ratio (11%), network stretch (10%), largest diameter (9%), node density (8%) and the node-surface distance (7%). These have in total 44% of the overall importance in the classification model (Fig. 6d). Interestingly, three of these five are nodal features of the network and two represent the overall morphology.

### 3.4. Mechanical Behavior Prediction based on Structural Features

The trained surrogate models predicted the mechanical response (simulation results) for small (2%) and large (20%) deformations purely based on structural features. Despite the high correlations between simulated and predicted mechanical parameters in both small and large deformations (Table 1), the performance metrics of the trained surrogate models increase with advancing compression (Fig. 7a-h). This specifically holds for predicting failure factors (*FR* in *α* = 0.02 → 0.20 : *R*^2^ = 0.96 → 0.97, *MPE* = 10% → 5% and *ŷ* = 0.95 → 1.03; *FB* in *α* = 0.02 → 0.20 : *R*^2^ = 0.81 → 0.95, *MPE* = 11% → 1% and *ŷ* = 0.91 → 1.00).

**Table 1:** Surrogate models performance in predicting calculated mechanical parameters in case of small (*α* = 0.02) and large deformations (*α* = 0.20).

| SM | $\alpha = 0.02$ | | | $\alpha = 0.2$ | | |
| --- | --- | --- | --- | --- | --- | --- |
| | Pred. $R^2$ | $MPE$ | Lin. fit $\hat{y}$ | Pred. $R^2$ | $MPE$ | Lin. fit $\hat{y}$ |
| $\bar{\sigma}$ | 0.90 | 4% | 1.04 | 0.96 | 5% | 0.97 |
| $\bar{\epsilon}$ | 0.89 | 14% | 1.11 | 0.93 | 9% | 0.99 |
| $FR$ | 0.96 | 10% | 0.95 | 0.97 | 5% | 1.03 |
| $FB$ | 0.81 | 11% | 0.91 | 0.95 | 1% | 1.00 |

**Figure 7:**
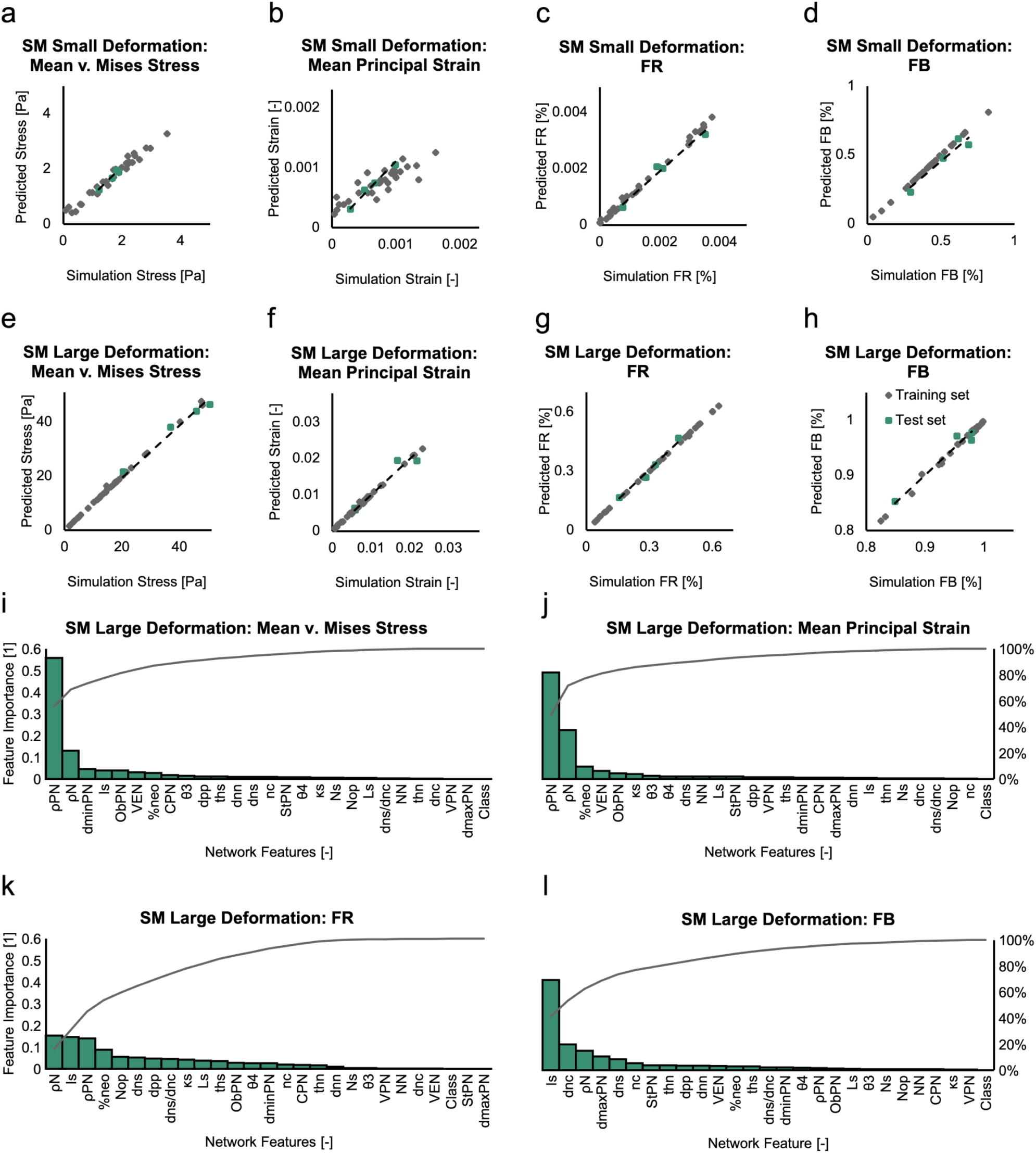
Surrogate mechanical models. a-d) Simulation results vs. surrogate model prediction for the test and training sets networks for small deformation in *EV* 3 primary direction. a) 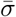. b) 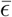. c) *FR*. d) *FB*. e-h) Simulation results vs. Surrogate model prediction for the test and training set networks for large deformation in *EV* 3 primary direction. e) 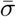. f) 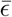. g) *FR*. h) *FB*. Training and test datasets are shown in gray and green, respectively. Dashed line represents a liner fit to the data points. i-l) Importance of structural features for the set of surrogate model predicting each mechanical parameter, i) 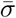. j) 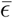. k) *FR*. l) *FB*. The gray line represents cumulative importance.

The structural features show different importance in predicting the mechanical parameters in case of large deformations. For both surrogate models predicting the mean stresses and strain, most important features (more than 70% of total importance) are network density (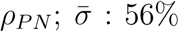 and 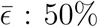) and node density (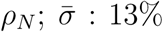 and 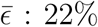; Fig. 7i, j). The prevailing structural features for predicting rupture failure factor are node density (*ρ*_*N*_ : 15%), segment inhomogeneity (*I*_*S*_: 15%), network density (*ρ*_*PN*_ : 14%) and percentage of open nodes (%*n*_*oe*_: 9%; Fig. 7k) with cumulative importance of more than 50%. In case of predicting buckling failure factor, segment inhomogeneity (*I*_*S*_: 41%), node-center distance (*d*_*nc*_: 12%) and node density (*ρ*_*PN*_ : 9%) are the most important features with cumulative importance of more than 60% (Fig. 7l).

## 4. Discussion

Structure and functionality of cytoskeletal protein networks are deeply linked. Although biochemical aspects have been thoroughly studied, little is known about the interplay between the structural characteristics of these networks with their mechanical functionality. Here we proposed a data-driven approach to investigate this structure-function relationship and presented the application to FtsZ protein network.

### 4.1. Mechanical Response of FtsZ Isoforms

We were able to show the mechanical response of a protein network to external loads and specifically that the precise structural response of FtsZ networks to compression is independent of the load direction and is different for FtsZ1-2 and FtsZ2-1 isoforms. This is to our knowledge the first detailed in silico investigation of the mechanical behaviour of cytoskeletal protein structures in response to external micro-environmental stimuli. This identified isoform-specific mechanical responses support the assumption of potentially different structural roles of these two main FtsZ isoforms [22]. Further, the isoform-specific mechanical responses are in accordance with the functional- and morphology-related observations of the same isoforms in yeast cells [19, 20].

In case of mechanical response of FtsZ in large deformation, we focus only on one direction due to four reasons: 1) the similar mechanical response at 2% displacement for all loading cases, 2) according to calculations in [23], variations of the two significant different parameters for all three principal directions (FtsZ1-2: 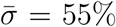 and FtsZ2-1: *FB* = 35%) can be explained by differences in stretch (FtsZ1-2 *St* = 0.76±0.11, FtsZ1-2 *St* = 0.67±0.20, *p* = 0.05), 3) the need of significant computational resources and 4) EV3 has the overall highest (combining FtsZ2-1 and FtsZ1-2) mean values at 2% plate displacement in all four parameters 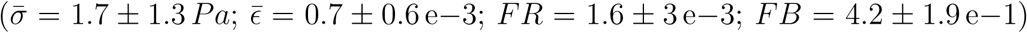. The similarity of mechanical responses between the two isoforms suggests that they contribute in response to large chloroplast deformation in a comparable (or combined) fashion to the plastid mechanics. Moreover, FtsZ isoforms show a semi-nonlinear increase in stress and strain with an increase in network deformation. This is similar to previously reported behavior of microtubule [54, 55] and actin filaments [56]. This points toward similar load-bearing functionality of FtsZ (as plastoskeleton). Furthermore, for compression up to 20%, buckling remains the prevailing failure factor. The convergence of *FB*, which reaches its limit at about 1%, suggests that the network minimizes the buckling probability, indicating an adaptive stability of FtsZ networks as previously suggested based on experimental observations [22, 57]. Although the rupture failure probability steadily increases with increasing compression, it remains significantly below the bulking failure factor, thus rupture failure is a less defining parameter for network failure. One reason for this might be, that FtsZ filaments experiencing high strain values leading to rupture only after buckling and at the location of bucking, which would be similar to the fragmentation of buckled actin filaments [58].

Previous studies employing simplified geometries, such as tensegrity models allowed theoretical studies of cellular mechanism such as cell reorientation [11, 15]. More detailed FE models have been developed to investigate the mechanical role of cytoskeletal components [13] and cell mechano-sensitivity [59]. However, the generic and strongly simplified geometrical representation of the cells, e.g ellipsoid [60, 61] (even when cytoskeleton filament directions were considered), potentially prevents comprehensive studies of the influence of structural features on the mechanical behavior of cytoskeletal protein networks. Our approach of performing nanoFE simulations on segmented 3D network geometries of life networks allows one to analyse structure-related aspects of protein network mechanics in a sample specific manner. Further, investigating the sub-cellular components separated from their surrounding allows to decouple protein network mechanics from whole cell mechanics. To date, contributions of cytoskeletal structures to whole cellular mechanics can only be indirectly inferred from experimental techniques such as AFM [62] and optical tweezers [63]. However, since the mechanical behavior of a cellular structure is determined by many components, such as of the structure of the cortical, intra-cellular (non-cortical) cytoskeletal, and nuclear networks, as well as their distribution in space, decoupling the individual components remains challenging [13, 64, 65]. It has been suggested that AFM measurements with sharp tips tend to emphasize biomechanical properties of the cell cortex, whereas AFM measurements with round-tips tend to emphasize stiffness of the intra-cellular network [65]. Combing such measurements with a structural detailed models, as shown here, would possibly further advance the understanding of cellular mechanics.

### 4.2. Structure-function Relationship in FtsZ Network

Our mechanical surrogate model is capable of predicting the mechanical behavior of protein networks in response to external loading. This holds for 2% as well as 20% compression (0.81 ≤ *R*^2^ ≤ 0.96 and 0.93 ≤ *R*^2^ ≤ 0.97, respectively). However, the relatively higher accuracy for large network compression points toward higher correspondence of the extracted structural features to the mechanical behavior of the network in response to large deformations. This specifically coheres to the hypothesis that these networks are able to undergo large deformations without losing their structural integrity, as previously postulated [22]; hence, possessing structural features conforming to response in case of large deformations. Moreover, the high accuracy in mapping the structural features to the mechanical behavior of the networks further demonstrates the potential load bearing functionality of FtsZ protein in chloroplasts. Furthermore, this shows that the capability of the network to keep its stability by undergoing deformations relies not only on material properties of the biopolymer, but probably more prominently, on the structural features of the network. This is in accordance with the effects of the network architecture on the overall mechanical behavior reported in actin protein network [66]. In summary, to our knowledge, this is the first detailed investigation of these sample-specific structure-based mechanical analysis of performance correlations. This enables us to not only have a image-based virtual mechanical testing method, but also a method to investigate the manifestation of the mechanical characteristics of structural network features.

By analyzing the importance of the features of the surrogate models in predicting stresses and strains of the network, we could show that the network density and the node density are the structural features mostly contributing to load bearing characteristic of the network. However, the mechanical failure behaviour of the networks is mostly corresponding to more local structural characteristics. Specifically, in the case of buckling failure behaviour, the surrogate model shows that local changes in the filament (segment inhomogeneity) and the distance of the nodes to the center dictate the observed mechanical behaviour. This can be interpreted as the network being capable of stopping the increase in failure possibility due to buckling of its filaments by possessing an arrangement of the nodes and filaments with the specific architecture that includes: local changes of direction and thickness of filaments in FtsZ1-2 and FtsZ2-1: *I*_*S*_ = 18.8 and the distance between the nodes and the center of the network: *d*_*nc*_ = 1.85 *µm*. This could potentially be used to design adaptively stable structures capable of undergoing large deformations [67] or mechanically optimized and synthetically engineered biomaterials [68, 69].

Our feature-based classification and regression models achieved on par accuracy with deep learning based protein network classification methods [70, 71], while adding the ability of extracting specific structural features enabling the predictions. This is specifically beneficial in the context of mechanical functionality, where our designed extracted features correspond to mechanical traits in their nature. Whereas, extracted features utilizing convolutional networks tend to be abstract and at best very challenging to interpret [72, 73]. Moreover, unlike deep learning models that require extremely large dataset for meaningful training, our models could reach high accuracy on *n* = 37 3D CLSM images.

### 4.3. Limitations

Our study has limitations. First, the imaging resolution might affect the simulation results as well as the mapping of the surrogate models. However, we have previously shown that our quantitative imaging method is capable of resolving the micro-structure of FtsZ networks [24]. Second, the commonly used linear elastic material model in FE simulations of cytoskeletons [13, 14, 59, 74] might not completely capture the mechanical behavior of the network. However, to our knowledge, to date no constitutive law has been developed for describing mechanical behavior of FtsZ. Although more complicated constitutive laws for the mechanical behavior of single actin filaments have been proposed [75–77], linear elasticity is the prevailing choice for cytoskeletal networks in whole-cell models [13, 14, 59, 74]. Moreover, our focus was not on exactly matching the mechanical behaviour, but on the influences of structural features. Future studies could focus on combining our approach of precisely modeling the micro-structure with experimental techniques, such as atomic force microscopy, to further investigate material properties of the FtsZ-based plastoskeleton. Third, the loading conditions of our simulations are not an exact duplication of reality, where a combination of active dynamic forces [40] as well as osmotic pressure [78] drive the morphological changes of the network. Moreover, the FtsZ isoform is surrounded by other proteins as well as other materials, such as inter-organelle fluids. Our designed simulation setup provides a generic platform to investigate the structure-function relationships in FtsZ protein network rather than a one-to-one simulation of dynamics of plastids. The failure criteria used in this study are experimentally derived from actin filaments [44, 79], since no failure criteria has been experimentally derived for FtsZ to date. However, due to the assumed similarity in structural functionality between the FtsZ network and actin networks and the similarity of rigidity in FtsZ and actin filaments, actin failure criteria might represent FtsZ behavior to a certain extent. Finally, the dataset size might restrict the generality of conclusions made by means of the introduced data-driven approach. Although the in-silico experimental setup increases the dataset size, more images would further improve the performance of the designed methodology.

### 4.4. Conclusions

In this work, we showed that combing confocal microscopy imaging with nanoFE analysis employing a machine learning framework allows for an image-based surrogate model capable of predicting cellular mechanical responses to external stimuli. Additionally, by providing a way to identify structural features determining the mechanical response with respect to a given stimulus, we were, for the first time, able to directly investigate the structure-function relationship of individual protein networks in a sample-specific manner. Our ML surrogate model trained on in-silico data generates highly accurate and fast predictions of isoform classification and the mechanical behavior on the sub-cellular level. Therefore, the method provides a framework to further investigate structural functionality of protein networks in plants as well as in humans, as it would allow to monitor the structure-function relationships of cytoskeletal components during morphological and, hence, time-dependent, changes, e.g., actin-driven cell shaping. This may also lead to an improved understanding of the mechanical aspects of cell-biomaterial interaction, and would provide insights into designing micro/nanoengineered functional biomaterials for future research on regenerative medicine, cell biology, development and diseases, as well as drug development.

## Acknowledgements

This study was supported by the Deutsche Forschungsgemeinschaft (DFG, German Research Foundation) under Germany’s Excellence Strategy – EXC 2075–390740016 (SimTech) and EXC 2189 (CIBSS), and as a part of Transregio/SFB TRR141, project A09.

## Conflict of interest statement

All authors declare no conflicting interests.

## Author’s roles

Conceptualization: PA, AB, RR, and OR. Contributing software tools: PA. Data processing and analysis: PA and AB, ZT. Molecular biology and microscopy: BÖ. Writing - original draft: PA and AB. Writing - review and editing: All. Funding acquisition: RR and OR.

## Notes

### Competing Interest Statement

The authors have declared no competing interest.

